# Out of the blue: the independent activity of sulfur-oxidizers and diatoms mediate the sudden color shift of a tropical river

**DOI:** 10.1101/2022.01.17.476333

**Authors:** Alejandro Arce-Rodríguez, Eduardo Libby, Erick Castellón, Roberto Avendaño, Juan Carlos Cambronero, Maribel Vargas, Dietmar H. Pieper, Stefan Bertilsson, Max Chavarría, Fernando Puente-Sánchez

**Affiliations:** Institute of Microbiology, Technical University of Braunschweig, D-38106, Braunschweig, Germany; Microbial Interactions and Processes Research Group, Helmholtz Centre for Infection Research, 38124, Braunschweig, Germany; Escuela de Química, Universidad de Costa Rica, 11501-2060, San José, Costa Rica; Centro de Investigación en Ciencia e Ingeniería de Materiales (CICIMA), Universidad de Costa Rica, 11501-2060, San José, Costa Rica; Centro Nacional de Innovaciones Biotecnológicas (CENIBiot), CeNAT-CONARE, 1174-1200, San José, Costa Rica; Centro de Investigaciones en Productos Naturales (CIPRONA), Universidad de Costa Rica, 11501-2060, San José, Costa Rica; Centro de Investigaciones en Estructuras Microscópicas (CIEMic), Universidad de Costa Rica, 11501-2060, San José, Costa Rica; Deparment of Aquatic Sciences and Assessment, Swedish University of Agricultural Sciences, Lennart Hjelms väg 9, 756 51, Uppsala, Sweden

**Keywords:** Geobiology, Hydroxyaluminosilicates, Hydrothermal, Sulfur Oxidizing Bacteria, Diatoms, Río Celeste

## Abstract

Río Celeste (“Sky-Blue River”) is a river located in the Tenorio National Park (Costa Rica) that has become an important hotspot for eco-tourism due to its striking sky-blue color. A previous study suggested that this color is not caused by dissolved chemical species, but by precipitation of light-scattering aluminosilicate particles at the mixing point of two colorless streams, the acidic Quebrada Agria and the neutral Río Buenavista. We now present microbiological information on Río Celeste and its two tributaries, as well as a more detailed characterization of the particles that occur at the mixing point. Our results overturn the previous belief that the light scattering particles are formed by the aggregation of smaller particles coming from Río Buenavista, and rather point to chemical formation of hydroxyaluminosilicate colloids with Quebrada Agria as the main contributor to the phenomenon, with Río Buenavista acting as a secondary source of silica to the reaction. We also show how the sky-blue color of Río Celeste arises from the tight interaction between chemical and biological processes. Sulfur-oxidizing bacteria generate an acidic environment in Quebrada Agria, which in turn cause dissolution and mobilization of aluminum and other metals, while in Río Buenavista the growth of diatoms transforms dissolved silicon into colloidal biogenic forms. The local interaction between these two well-known biological activities gives rise to the unique Río Celeste phenomenon, in what constitutes a textbook example of emergent behavior in environmental microbiology.

## 1. Introduction

Costa Rica is located in the Pacific rim of fire and therefore has a number of active volcanoes, and multiple manifestations of hydrothermal and volcanic origin such as thermal springs, acidic rivers or mineral-rich streams (Arce-Rodríguez *et al*., 2017, 2019). One of the most striking manifestations of that activity, that can be observed in the complex basaltic-andesitic volcanic massif of Tenorio (Guanacaste Volcanic Mountain Range), is Río Celeste (“Sky-Blue River”). Located within the Tenorio Volcano National Park, this river is recognized as one of the most beautiful rivers in the world for its characteristic sky-blue color that contrasts with the dark surrounding rainforest (**Figure 1**). Because of this, the river and the landscapes of the National Park have become an important hotspot for eco-tourism in Costa Rica (Jovanelly *et al*., 2017), attracting more than 100,000 visitors per year (SINAC, 2019).

**Figure 1.**
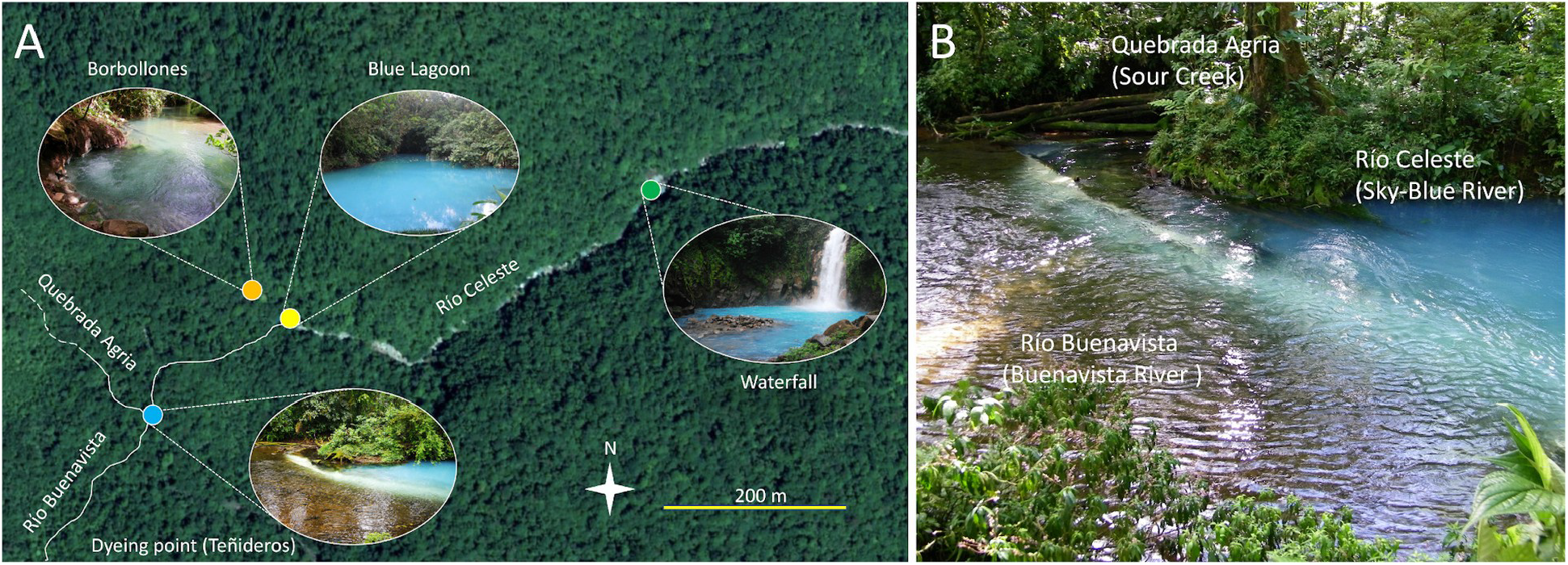
Environmental setting. **a)** Satellite view of the Río Celeste area. River paths are shown with continuous lines when they are not directly evident from the satellite imagery. Dashed lines in Quebrada Agria indicate that the upstream of this river is still uncharted. **b)** Detail of the ‘Teñidero’ (‘dyeing point’) confluence, showing the confluence between Río Buenavista and Quebrada, and the origin of the sky-blue river Río Celeste.

What makes Rio Celeste unique is that it is possible to observe the specific site where the river turns blue. At this point known locally as ‘Teñidero’ (dyeing point), the two colorless streams Río Buenavista (Buenavista River) and Quebrada Agria (Sour Creek) merge, and the resulting waters quickly acquire the sky-blue color (**Figure 1**). In a previous study, (Castellón *et al*., 2013) showed that the sky-blue coloration of Río Celeste is caused by Mie light-scattering caused by particles suspended in its water stream. This phenomenon has been scarcely studied (Ohsawa *et al*., 2002; Oyama & Shibahara, 2009, Takagai *et al*., 2016) and remains poorly understood. Because our previous measurements indicated that the particles present in Río Buenavista were smaller than the particles present in Río Celeste after the ‘Teñidero’ confluence (Castellón *et al*., 2013), we initially proposed that the light-scattering particles originated via a pH-dependent aggregation mechanism when Río Buenavista (pH=6.8) becomes acidified by Quebrada Agria (pH=3.1), marking the beginning of Río Celeste (pH=5.5). At that point we did not have measurements on the composition of the suspended particles, and we therefore alternatively examined the white material conspicuously deposited on the rocks which showed properties consistent with the presence of an aluminosilicate mineral.

While our previous work demonstrated the role of light-dispersing aluminosilicates in giving Río Celeste its characteristic color, the formation mechanism of such particles was however not elucidated. Furthermore, our original study did not take into consideration the potential role of microorganisms in generating the chemical conditions conducive to this process. In the present study we provide a detailed analysis of the colloidal-sized particles that occur at the ‘Teñidero’ dyeing point, and examine the microbial communities that inhabit Rio Buenavista, Quebrada Agria and the combined microbial community in Río Celeste. Our results challenge the previous perception in favor of a model where the particles causing Mie scattering mainly originate from dissolved aluminum and silicic acid coming from Quebrada Agria, with Río Buenavista acting as a secondary source of silica. Microbial community analyses show that Quebrada Agria hosts sulfur-oxidizing bacteria that are likely responsible for the acidic conditions in that source water, indirectly promoting aluminium mobilization. In contrast, Rio Buenavista is a source of diatoms to Río Celeste whose presence influences silicon speciation and partitioning. Both the presence of sulfuroxidizing bacteria in Quebrada Agria and the presence of diatoms in Río Buenavista are consistent with the chemical conditions of the respective stream. Our results shed new light on the combined biological and chemical processes that interactively cause formation of aluminosilicate particles that produce the beautiful sky-blue color that Río Celeste is well known for.

## 2. Materials and methods

### 2.1. Site description

The ‘Teñidero’ confluence (10°42’02.5” N, 84°59’49.3” W) is located in the Costa Rican Tenorio Volcano National Park, at the confluence of two streams: the neutral Río Buenavista and the acidic Quebrada Agria (Sour Creek). The whole area is embedded in the Volcanic Mountain Range of Guanacaste, and the region experiences frequent hydrothermal activity which give rise to a multitude of niches of volcanic origin such as thermal waters, acidic rivers or hydrothermal springs (Arce-Rodriguez et al. 2019). Even though some other rivers show some sky-blue color, the ‘Teñidero’ stands out as a unique point of research and popular interest since only here the confluence of two seemingly colorless streams instantly turn into the sky blue-colored Río Celeste (**Figure 1**).

### 2.2. Sampling and field measurements

All necessary permits for sampling water and sediments were obtained from the National System of Conservation Areas (SINAC) of the Ministry of Environment and Energy (MINAE) of Costa Rica (Resolution No. 097-2014-ACAT). On 19 January 2014, samples of water and sediments were collected from Rio Buenavista and Quebrada Agria about 5 meters before their confluence (10°42’02.5” N, 84°59’49.3” W). Rio Celeste’s samples were taken 20 meters after the confluence. At each sampling point, 3 water samples (1L each) and 3 sediment samples (between 10-50 g) were taken across the river. All samples for DNA analysis were collected in clean and sterile glass bottles, chilled on ice, stored at 4 °C, pooled and processed within less than 24 h. For qualitative analysis of diatoms, three water samples (50 mL) were collected in 50-mL Falcon tubes for each stream. Biofilm samples were also collected by brushing the surface of stones from the bottom of Río Celeste. All samples were fixed with 1% lugol’s iodine solution and preserved with 3% neutralized formaldehyde. The chemical and physicochemical parameters for the three streams can be found at **Supplementary Table S1 (Castellón et al. 2013)**.

### 2.3. Characterization of suspended materials

Water samples from Quebrada Agria, Río Buenavista and Río Celeste (1 L each) were individually filtered through porous membranes with a nominal pore size of 0.4 μm. The collected filters were allowed to dry at room temperature (20-25 °C). No residues were observed on the filter from Quebrada Agria. The residues collected from the water samples of Río Celeste and Río Buenavista were adhered to electrically conductive carbon tape for their analysis on a scanning electron microscope (SEM, Jeol JSM-IT500). This apparatus was equipped with a detector for energy dispersive x-ray spectroscopy (EDS) to determine the bulk chemical composition of the suspended materials collected in the filters.

For the precipitation experiments, water samples from Río Buenavista and Quebrada Agria (100 mL each) were respectively acidified with 0.005 M H_2_SO_4_ and neutralized with 0.01 M Na_2_CO_3_ until they reached the pH of Río Celeste (pH 5.5). The neutralization of the sample from Quebrada Agria produced suspended particles that were collected by centrifugation. The collected particles were resuspended in de-ionised water, centrifugated again and subsequently dried at room temperature. Such particles were pasted on carbon tape and analysed by SEM-EDS.

### 2.4. Total DNA isolation, construction of 16S rRNA gene libraries and Illumina sequencing

Three water samples (1L each) taken at different points along the river width were filtered through a vacuum filtration system under sterile conditions using a membrane filter (pore size 0.22 μm; Millipore, GV CAT No GVWP04700). To prevent filter rupture, a support membrane (pore size 0.45 μm; Phenex, Nylon Part No AF0-0504) was placed below. The upper filter was collected and stored at −80 °C until processing. The DNA was extracted from aseptically cut pieces of the filter using the PowerSoil^®^ DNA Isolation Kit (MoBio, Carlsbad, CA, USA) as described by the manufacturer. Cells were disrupted by two steps of bead beating (FastPrep-24, MP Biomedicals, Santa Ana, CA, USA) for 30 s at 5.5 m s^-1^. For the construction of microbial 16S rRNA amplicon libraries, the V5-V6 hypervariable region were PCR-amplified with universal primers 807F and 1050R (Bohorquez et al. 2012). Barcoding of the amplicons and addition of Illumina adaptors were conducted by PCR as described previously (Camarinha-Silva et al. 2014; Burbach et al. 2016). The PCR-generated amplicon libraries were subjected to paired-end 2×250 Illumina MiSeq sequencing (Illumina, San Diego, CA, USA).

### 2.5. Bioinformatic analysis of 16S rDNA amplicon data

Raw MiSeq sequences were quality-filtered and merged with *moira.py* v1.3.2 (Puente-Sánchez et al. 2016) with options *--paired --ambigs disallow --consensus_qscore posterior --qscore_cap 0 --maxerrors 1 --collapse True --alpha 0.005 --output_format fastq*. A custom *python* script was used to expand the derreplicated files generated by *moira.py* so they could be used in subsequent analysis steps, and the contigs were then derreplicated and chimera screened with DADA2 v1.14.1 (Callahan *et al*., 2016). The resulting Amplicon Sequence Variants (ASVs) were aligned to the SILVA nr 132 reference database (Quast *et al*., 2012) using *mothur v1.44.0* (Schloss *et al*., 2009). The alignment was curated with *mothur* using the *screen.seqs* (options: *start=25318, end=33595, maxhomop=7, minlength=200, maxlength=275*) and *filter.seqs* (options: *vertical=T, trump=.*) commands, after which a UPGMA tree was constructed from the filtered alignment using *phangorn* v2.5.5 (Schliep, 2011). Variance-adjusted weighted UniFrac distances (Lozupone *et al*., 2007) were obtained from the tree with *GUniFrac* v1.1 (Chen & Chen, 2018). Following McMurdie & Holmes (2014), sequences were not rarefied prior to weighted UniFrac distance calculation. These distances were in turn used to compute a Non-metric Multidimensional Scaling with the *metaMDS* function from *vegan* v2.5.6 (Oksanen *et al*. 2013). Further, we performed permutational multivariate analysis of variance (Chen *et al*., 2012) as implemented in the *PermanovaG* function from *GUniFrac* package to assess whether community composition was influenced by sample type (Water vs Sediment) and origin (Hydrothermally influenced vs Neutral). Finally, we used SINA (Pruesse *et al*., 2012) to taxonomically classify the ASV sequences, and SQMtools (Puente-Sánchez *et al*., 2020) to generate barplot figures. The following identity cutoffs to the closest match in the database were assigned to assign ASVs to the different taxonomic ranks (Yarza *et al*., 2014): phylum=75%, class=78.5%, order=82%, family=86.5%, genus=94.5%. The taxonomy of each ASV is thus reported at the maximum resolution attainable without compromising a correct classification. Raw sequences were submitted to the sequence-read archive (SRA) under BioProject PRJNA747923. The sequencing data from the hydrothermal spring Borbollones (located also inside the Tenorio Volcano National Park; Figure 1) were obtained from our previous study (Arce-Rodríguez *et al*., 2019).

### 2.6. Identification of diatoms by optical and electronic microsocopy

Both the water samples and biofilms were sedimented at room temperature for 24 hours. The sedimented material was examined under a light microscope (Model IX-51, Olympus) at 40X or 100X optical magnification. Selected samples were analyzed in a scanning electron microscope (Model S-3700N, Hitachi) using an accelerating voltage of 15 kV.

## 3. Results

### 3.1. Formation of minerals at the ‘Teñidero’ confluence

When acidic Quebrada Agria flows into Río Buenavista, the well-defined zone of mineral precipitation is in near steady-state and is known locally as the ‘Teñidero’ or dyeing point. Some of the material eventually precipitates as white deposits on the river rocks but in the present study we focus on the light-scattering particles that remain in suspension. According to the dynamic light scattering (DLS) measurements taken in Castellón (2013), these particles have an average diameter of 566 nm, much larger than the 184 nm particles that are suspended in Río Buenavista. This leads to preferential Mie scattering of shorter wavelengths (e.g. blue and violet solar radiation) responsible for the sky-blue color of Río Celeste. In order to determine the nature and origin of the light-scattering particles, we isolated the suspended material from the streams of Río Buenavista and Río Celeste, and analyzed them in bulk with scanning electron microscopy (SEM, **Figure 2**), energy dispersive X-ray spectroscopy (EDS, **Supplementary Figure S1**) and Fourier transform infrared spectroscopy (FTIR, **Supplementary Figure S2**). Quebrada Agria lacked relevant suspended particles.

**Figure 2.**
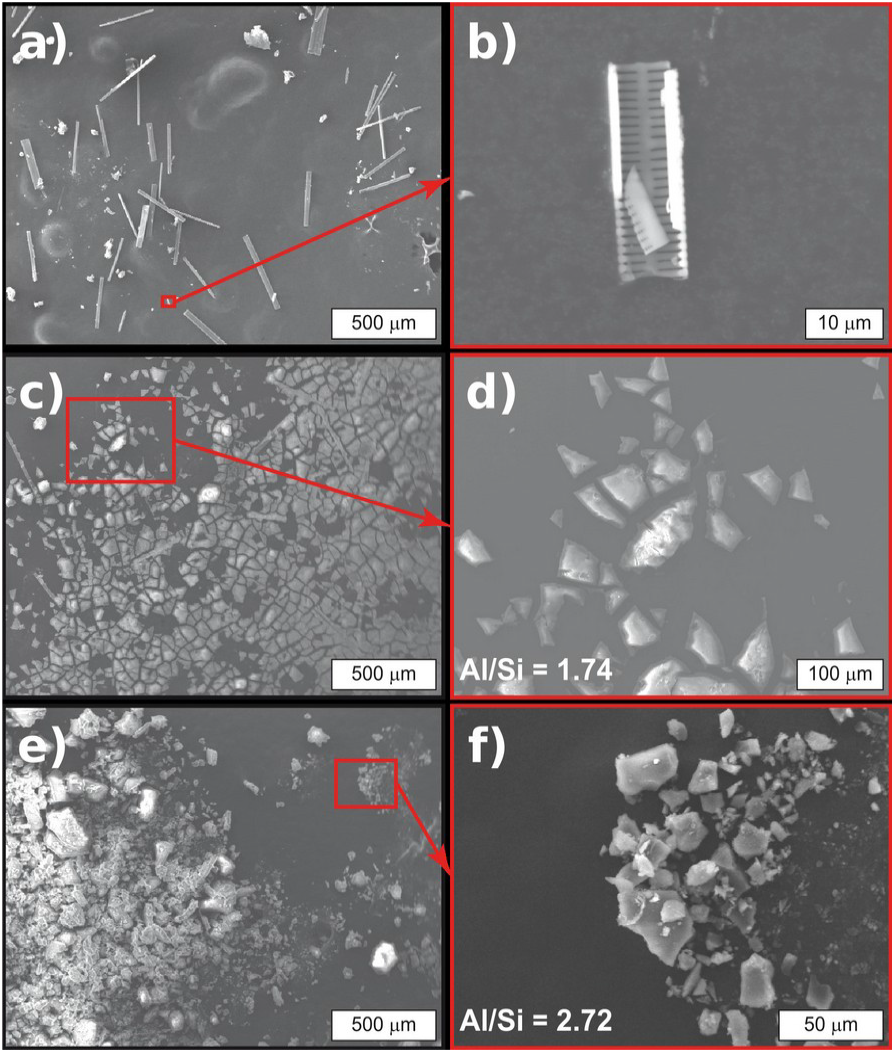
Scanning electron microscopy (SEM) images of filtered or precipitated materials from the water streams at the confluence that originate Río Celeste. The atomic Al/Si ratios were obtained by EDS scans. **a)** Filtered materials from Río Buenavista; the solids retained by the membranes are mainly frustule fragments. **b)** Detail of a frustule fragment. **c)** Filtered materials from Río Celeste; frustule fragments along with agglomerated particles are retained by the filter membranes. **d)** Magnification showing a detail of the agglomerated filtered particles from Río Celeste. **e)** Agglomerated particulate materials obtained by *in vitro* precipitation of a water sample from Quebrada Agria. **f)** Magnification performed on the specimen in **e)**, showing the agglomerated particles.

In agreement with our previous study, suspended matter in Río Buenavista consisted of particles that were too small to effectively scatter light according to the Mie principle, mixed with a few larger particles (e.g. frustule fragments) (**Figure 2a,b**). In stark contrast, we found abundant particles in Río Celeste River water that formed larger aggregates in the filtered sample (**Figure 2c,d**). The fact the aggregates were larger than the diameters previously predicted by Dynamic light scattering (DLS) can be explained by flocculation during filtering, drying and centrifugation, as it has been described previously (Comba & Kaiser, 1990), as well as by the removal of the smaller-sized particles during filtering. In spite of this discrepancy, we find it safe to assume that the aggregates were representative of the light-scattering suspended materials. A composition map assessed on a broad region (11.0 μm x 8.3 μm) of the solid from Río Celeste indicated the presence of aluminum, silicon and oxygen together with small and variable amounts of sulfur (likely adventitious sulfate from the water), with an Al/Si ratio of 1.74. Fourier-transform infrared (FTIR) spectroscopy revealed a structure typical of a silicon-rich hydroaluminosilicate (**Supplementary Figure S2**; Lenhardt *et al*., 2021).

To verify the mechanism of particle precipitation, a new *in vitro* experiment was performed. Samples of water from the neutral Río Buenavista and the acidic Quebrada Agria were respectively acidified and neutralized, until both reached the pH of Río Celeste. Acidification of Río Buenavista water did not produce any particles while the elevation of pH in Quebrada Agria waters produced abundant particles rich in aluminum, silicon and oxygen, with an Al/Si ratio of 2.72 (**Figure 2e,f**).

### 3.2. Microbial community composition at the Quebrada Agria (QA) stream

The microbial community in Quebrada Agria was dominated by Proteobacteria and Campylobacterota (**Figure 3a**, QAW) assigned to known sulfur-oxidizing genera such as *Sulfuriferula* (**Figure 4**, ASVs 1, 18), *Halothiobacillus* (ASV 3), *Sulfurimonas* (ASVs 4, 5), *Thiomonas* (ASV 44) or *Acidithiobacillus* (ASV 46). Archaea (Thaumarchaeota and Crenarchaeota) were also detected. Overall, the microbial composition of Quebrada Agria waters resembles to that of Borbollones (**Figure 3a**, BBW, see hierarchical clustering), a nearby-located (at *ca* 300 m) hydrothermal spring whose microbial composition was recently described (Arce-Rodríguez *et al*., 2019). Overall, the sediment featured similar ASVs as the waters, but in different proportions (**Figure 3a**, QAS). Campylobacterota were missing and Proteobacteria were present in lesser amounts and dominated by the iron-oxidizing family Gallionellaceae (**Figure 4**, ASVs 14, 24), while the proportion of Archaea (Nitrosotaleaceae family, ASV 7, 26), Actinobacteria, Acidobacteria, Nitrospirota and Planctomycetota were higher. Some high-abundance ASVs were endemic to the QA sediments; this includes an Acidothermaceae actinobacterium (ASV 47) and a Nitrosomonadaceae gammaproteobacterium (ASV 81).

**Figure 3.**
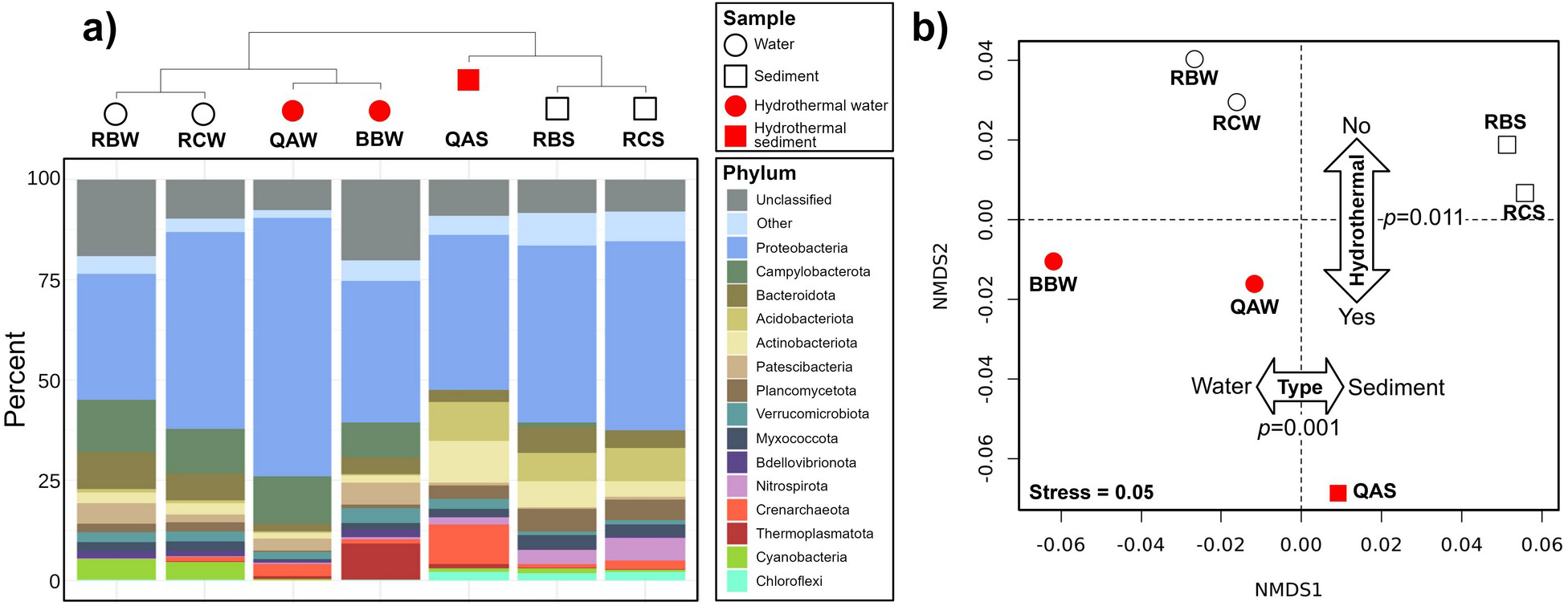
Microbial community composition around the ‘Teñidero’ confluence. Samples were from the two streams converging into the dyeing point (sample codes starting by RB and QA) and the resulting mixed stream (‘Río Celeste’, sample codes starting by RC). For each location, we analyzed plankton from the water (W) and sediments (S). An extra water sample from a nearby hydrothermal spring (BBW) was also included for comparative purposes. **a)** Hierarchical clustering of the samples based on their variance-adjusted weighted UniFrac distances (top) and barplot showing their phylum-level composition (bottom). **b)** Non-metric multidimensional scaling (NMDS) of the samples included in this study. Plotting symbol and color denote sample type (circle: water, square: sediment) and origin (white: neutral samples; red: hydrothermally-influenced samples). Two-way arrows and the associated p-values indicates a significant influence (PERMANOVA test using UniFrac distances) of sample type and origin in microbial community composition.

**Figure 4.**
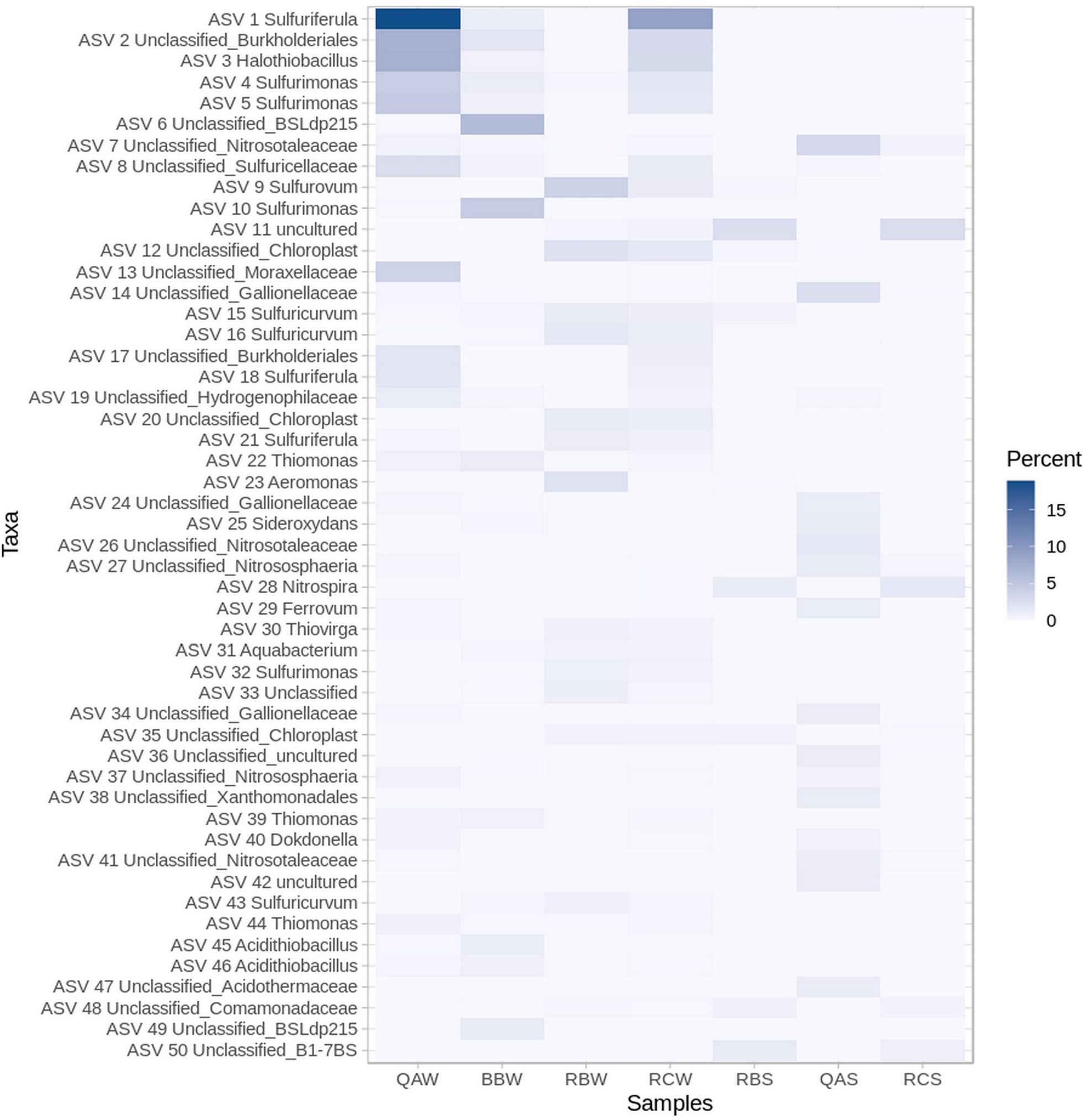
Distribution and taxonomy of the 50 most abundant ASVs. Heatmap intensity represents percentage abundance. The first two letters in the sample code indicate the river (QA: Quebrada Agria, BB: Borbollones, RB: Río Buenavista, RC: Río Celeste), the third letter indicates the material (W: water, S: sediment).

### 3.3. Microbial community composition at the Río Buenavista (RB) stream

The most abundant phyla in Buenavista were Proteobacteria, Campylobacteriota and Bacteroidota (**Figure 3a**, RBW). Among them, the most abundant bacterial genus was *Sulfurovum* (**Figure 4**, ASV 9) followed by *Aeromonas* (**Figure 4**, ASV 23). Many of the abundant bacterial taxa were related to the recycling of sulfur and nitrogen, such as *Sulfurimonas* (ASV 32), *Thiovirga* (ASVs 30), *Sulfuricurvum* (ASVs 43), *Thiotrix* (**Supplementary Table S2**, ASV 53), Galliollenaceae (ASV 130) and Comamonadaceae (ASV 149). For eukaryotes, the second most abundant ASV in Buenavista’s waters was a chloroplast (**Figure 4**, ASV 12), and in general chloroplasts represented the majority of the cyanobacteria-related reads obtained in this study (**Supplementary Figure S3**). Since the results obtained with SINA does not distinguish between different chloroplasts, we manually annotated the sequence of ASV12 using the online Nucleotide BLAST tool from NCBI (Boratyn *et al*., 2013). The first hit was a 100% match to the chloroplast of the Bacillariophyceae diatom *Halamphora coffeaeformis* (NCBI accession number NC_044465.1), but several other perfect matches to other diatom species were also found, implying that the V5-V6 16S rRNA region amplified in this study is not good at discriminating between different diatom groups. In order to achieve a better classification, we instead resorted to optical and electronic microscopy (see section 3.5). As for Quebrada Agria, the microbial population in Buenavista’s sediment was a mixture of endemic taxa and taxa which were also present in the water. Campylobacterota were at much lower relative abundances than in the water, and Actinobacteria, Proteobacteria, Acidobacteria, Nitrospirota and Planctomycetota increased (**Figure 3a**, RBS). Among the taxa present in the sediments but not in the waters, the most abundant belonged to the Alphaproteobacteria and Gammaproteobacteria classes (**Supplementary Table S2**, ASVs 50, 58, 66, 86, 88) and the Bacteroidota and Nitrospirota phyla (ASV 185 and 120, respectively).

### 3.4. Microbial community composition at the Río Celeste (RC) stream

The composition of Río Celeste’s planktonic microbiome was overall more similar to that of Río Buenavista than to that of Quebrada Agria (**Figure 3b**), which is to be expected since Buenavista is the largest source of Celeste’s waters (*ca*. a 3:1 proportion with Quebrada Agria according to mass balance of conservative elements). The contribution of Quebrada Agria was still apparent from the high relative abundance of Proteobacteria (*Sulfuriferula*; **Figure 4**, ASVs 1, 18) and Archaea. The sediment of Río Celeste was very similar to that of Río Buenavista, except for the higher presence of a few ASVs (**Figure 4**, ASV 7 – Nitrosotaleaceae; **Supplementary Table S2**, ASVs 61 – Nitrospira and 74 - Uncultured Planctomycetales) attributed to the influence of Quebrada Agria.

### 3.5. Diatom composition at the Río Celeste (RC) stream

The analysis by optical- and electron-microscopy validated the results obtained by sequencing. A great variety of diatoms with different morphologies and sizes between 2-200 μm were observed (**Supplementary Figure S3**). Morphologically, we identified organisms from the Bacillariophyta and Miozoa phyla. The identified species can be grouped into 3 classes (Bacillariophyceae, Mediophyceae and Dinophyceae) and 6 families (Diploneidaceae, Biddulphiaceae, Surirellaceae, Cymbellaceae, Catenulaceae and Peridiniaceae). Specifically, diatoms of genera *Diploneis, Biddulphia, Surirella, Cynbella, Amphora*. The dinoflagellate *Peridinium* was also recorded being one of the most abundant taxa observed with light microscopy (**Supplementary Figure S3b**).

## 4. Discussion

### 4.1. Quebrada Agria (QA) is influenced by upstream hydrothermal activity

Quebrada Agria is a transparent and acidic stream (pH 3.1) largely devoid of suspended materials (Castellón *et al*., 2013) while having a high content of dissolved sulfate (190 mg/L), chloride (71 mg/L), calcium (55 mg/L), and aluminium (12 mg/L) (**Supplementary Table S1;** Castellón et al. 2013). While this distinctive chemistry suggests the influence of hydrothermal activity, the source of Quebrada Agria’s waters could not be determined during our sampling campaign and likely lies within the Tenorio Park rainforest. We therefore turned to the microbial community composition, and found that it was similar to that of the nearby (*ca* 300 m) Borbollones hydrothermal spring (Arce-Rodríguez *et al*., 2019; sample code BBW). Samples from Quebrada Agria and Borbollones had a significantly different composition compared to the rest of the samples (**Figure 3b**, PERMANOVA *p* = 0.011), and both shared prominent and functionally defined microbial taxa such as the sulfur-oxidizers *Sulfurimonas* and *Sulfuriferula* and the iron-oxidizer *Gallionela* (**Supplementary Table S2**). These genera have also been found in other hydrothermally influenced settings (Meier *et al*., 2017; Lopez-Bedogni *et al*., 2020; Li *et al*., 2012).

Based on these chemical and microbiological similarities, we hypothesize that Quebrada Agria is influenced by one or more upstream hydrothermal springs akin to the one described in Arce-Rodríguez *et al*., 2019). Under this scenario, sulfide originating from hydrothermal fluids would be oxidized by microbial activity (Amils *et al*., 2007; Arce-Rodríguez *et al*., 2020), causing a decrease in pH and high sulfate concentrations observed near the ‘Teñidero’ site. The acidity of the water would in turn promote weathering and result in extensive mobilization of dissolved silicon and aluminum (Bigham & Nordstrom, 2000).

### 4.2. Río Buenavista (RB) is a circumneutral stream with standard chemical composition

The Río Buenavista has many of the features typically seen in freshwater ecosystems: pH is close to neutral (6.8) and the chemical composition of the water is within the established levels of metals and anions suitable for human consumption (**Supplementary Table S1**). The river is largely transparent and shows no noticeable suspended particles but do contain ample amorphous colloidal silica particles with an average diameter of 184 nm (Castellón *et al*., 2013). The most abundant ASV in Buenavista’s waters were assigned to the *Sulfurovum* genus (**Figure 4**, ASV 9). Other genera related to sulfur cycling such as *Sulfurimonas, Thiovirga* or *Thiotrix* were also present in moderate amounts. The dominance of *Sulfurovum* is somewhat puzzling, as this genus is usually found in hydrothermal vents (Meier *et al*., 2017), cold seeps (Niemann *et al*., 2013), groundwater aquifers (Hubalek *et al*., 2016), freshwater sediments (Duan *et al*., 2020) or as endosymbionts of larger organisms (Salonen *et al*., 2019). It is unclear if any of these conditions would apply in Río Buenavista waters: they could be originating from the sediments (where they are also present, albeit at lower relative abundance), be indicative of hydrothermal activity (although Buenavista’s waters have an apparently standard chemical composition), or originate from some nearby groundwater seep.

The second most abundant bacteria (third most abundant ASV including non-bacterial sequences) was an ASV from the heterotrophic *Aeromonas* genus (**Figure 4**, ASV 23). *Aeromonas* is commonly seen in freshwaters and drinking waters (Holmes *et al*., 1996), and appears either as a free-living organism or associated to crustaceans or fish, for which they can act as pathogens (Janda & Abbot, 2010). The abundance of this genus in freshwaters has been considered an indicator of trophic status, analogous to Secchi disc depth, total phosphorus, or chlorophyll *a* (Rippey & Cabelli, 1989). Their high relative abundance suggest that Río Buenavista is somewhat eutrophic.

Finally, the second most abundant ASV in Río Buenavista’s waters was classified as a Chloroplast (**Figure 4**, ASV 12). Manual search against the NCBI *nr* database indicated that the chloroplast 16S rRNA gene sequence is from a diatom but could be affiliated to several taxonomic groups (**Supplementary Table S3**). This was confirmed by light and electron microscopy, which revealed the presence of the diatom genera *Diploneis, Biddulphia, Surirella, Cynbella* and *Amphora* (**Supplementary Figure S3**). We note that the abundance of diatoms in Río Buenavista might be much higher than suggested by the relative abundance of ASV 12, since our sample processing protocol was not optimized for retrieving DNA from hard-to-lyse eukaryotic cells. Our observations are nonetheless consistent with previous reports of high diatom activity in tropical rivers with low turbidity (De Master *et al*., 1983).

### 4.3. Quebrada Agria is the main contributor to the generation of light-dispersing colloidal hydroxyaluminosilicates

In our previous article (Castellón *et al*., 2013) we proposed that the light-dispersing particles from Río Celeste formed upon the aggregation of smaller colloidal aluminosilicate particles coming from Río Buenavista. In this work, while examining the particulate materials in water from Río Celeste, we observed an abundance of amorphous particles large enough to prompt us to question our original hypothesis. To this end we analyzed the suspended material that clearly originated at the ‘Teñidero’ confluence by SEM/EDX (**Figure 2b**). The particle morphology, abundance and composition obtained in these analyses strongly suggested that a chemical precipitation reaction was taking place at the mixing point. The Fourier-transform infrared (FTIR) spectrum of the Río Celeste particles was similar to that observed for short-range ordered aluminosilicates (SROAS) obtained under ambient conditions (compare **Supplementary Figure S1** with Figure 4 in Lenhardt *et al*., 2021), suggesting that a similar precipitation mechanism might be in place. The Al/Si ratio of these substances, which are intermediates in the formation of other minerals, varies between 2 and 1 and are also named hydroxyaluminosilicates HAS_A_ and HAS_B_ respectively. (Beardmore 2016).

To further explore the origin of these particles, we performed an *in vitro* experiment in which water samples from the neutral Río Buenavista and the acidic Quebrada Agria were respectively acidified and neutralized. The acidification of the water sample from Río Buenavista produced no noticeable changes upon reaching the pH of Río Celeste. However, a conspicuous proliferation of colloids was observed on the sample from Quebrada Agria upon its neutralization (**Figure 2c**). This strongly suggests that, contrary to our original hypothesis, material from Quebrada Agria is the main source of the Río Celeste particles. The Al/Si atomic ratio in the synthetic particles was 2.72, higher than the 1.74 observed in the naturally-occurring particles in Río Celeste, suggesting that Río Buenavista played a secondary role by providing extra silicon to the reaction. As the particles from Río Buenavista detected by dynamic light scattering in the first study had diameters that were on average 184 nm (Castellón *et al*., 2013), it can be assumed that these smaller particles become embedded with the particulate material formed upon reaction with aluminum and silicic acid from Quebrada Agria.

### 4.4. Diatoms as sources of colloidal silica in Río Buenavista

Diatoms incorporate dissolved silicate into their frustules, being the main source of biogenic silica in riverine settings (Conley, 1997; Ran *et al*., 2016) and under some circumstances accounting for the majority of the total silicon budget (Admiraal *et al*., 1990; Conley 1997). Most of the biogenic silica in rivers is not actually associated to living diatoms, but to detrital particles that originate when diatoms die or are preyed upon (Krause *et al*., 2010; Carbonell *et al*., 2013). Interestingly, diatom cultures have been shown to produce extracellular silica nanoparticles in the 150-400 nm diameter range (Losic *et al*., 2008), which is compatible with the 184 nm average diameter of the colloidal silica particles found in Río Buenavista (Castellón *et al*., 2013). Lithogenic silica (coming from the weathering of minerals such as clays, silts and sand) can also represent a large fraction of the total silicon in rivers (Huang *et al*., 2019). The particle size from such processes is however in the micrometer scale (Kravchishina *et al*., 2011; Le Meur *et al*., 2016), much larger than the particles detected here (Castellón *et al*., 2013). Additionally, Río Buenavista was transparent and devoid of apparent particulate material. Overall, the data are consistent with the notion that diatoms, rather than lithogenic sources, are responsible for the colloidal silica present in the water from Rio Buenavista. Diatom activity would therefore control the partitioning of silicon between the dissolved and colloidal forms in Río Buenavista, making them indirect contributors to the Río Celeste phenomenon.

### 4.5. Biological influence in the distinctive color of Río Celeste

Río Celeste arises at the ‘Teñidero’ confluence, when the acidic and aluminum-rich Quebrada Agria mixes with the neutral Río Buenavista. Our experiments suggest that the neutralization of Quebrada Agria initiates a precipitation reaction in which aluminum and silicon precipitate with dissolved and colloidal silica coming from Río Buenavista, resulting in colloidal hydroxyaluminosilicates that scatter light as described in Castellón *et al*., 2013. The interaction of two streams with a characteristic physicochemical composition is thus necessary for the emergence of the unique Río Celeste phenomenon. Our data strongly suggest that, in both streams, this composition is biologically mediated (**Figure 5**).

**Figure 5.**
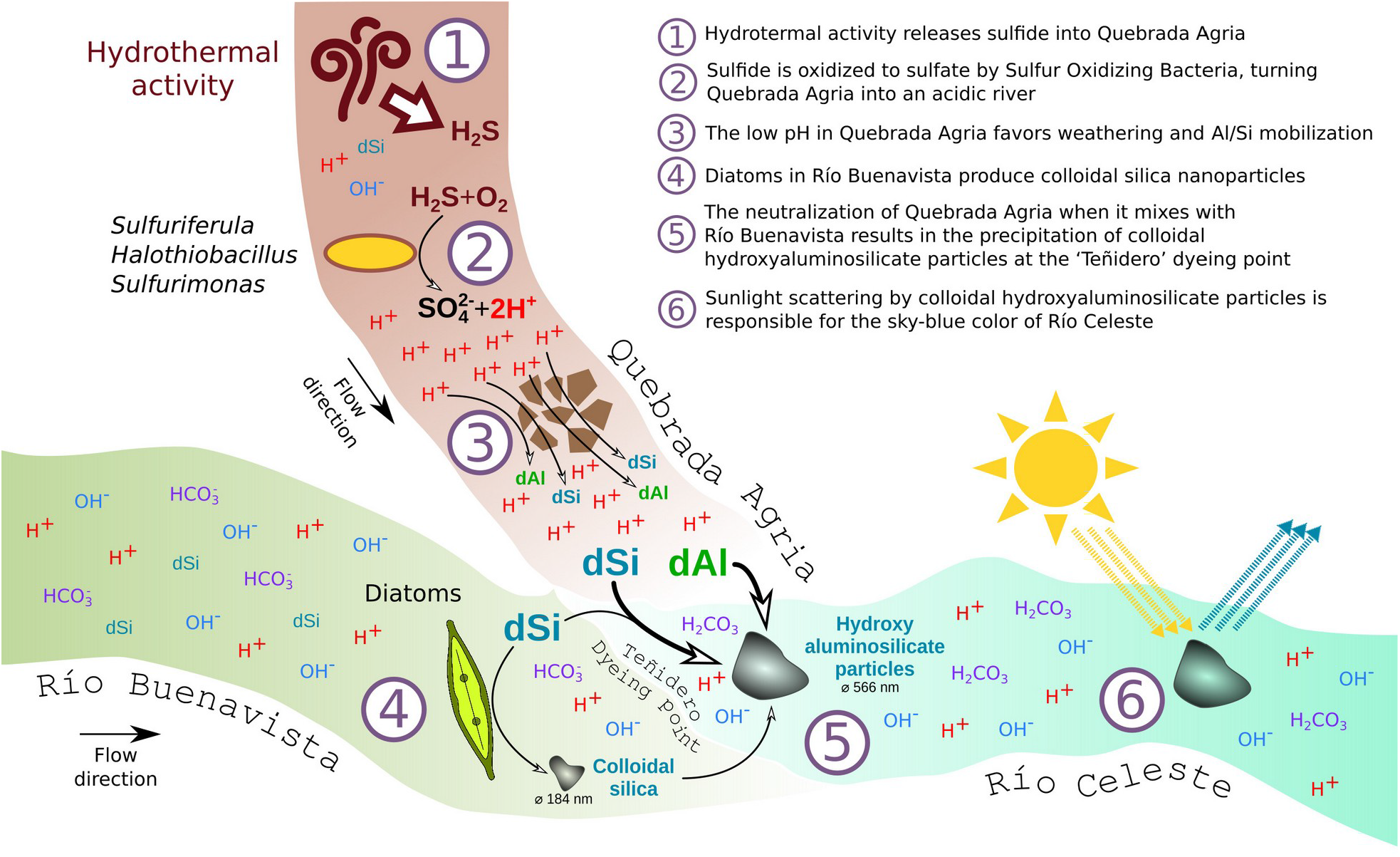
Conceptual biogeochemical model of the origin of Río Celeste’s sky-blue color.

In Quebrada Agria, hydrothermal sulfide is oxidized by bacteria such as *Sulfuriferula and Sulfurimonas* resulting in an acidic fluid capable of mobilizing silicates and metals such as aluminum, as has been described for other ecosystems (Bigham & Nordstrom, 2000; Amils *et al*., 2007; Arce-Rodríguez *et al*., 2020). Meanwhile, the partitioning of silicon in Río Buenavista is controlled by diatoms, which are in turn the most likely source for the colloidal silica particles suspended in the stream. Diatom particles are known for their ability to act as flocculants (Passow & Alldredge, 1995; Hamn, 2002), and incorporate aluminium both during biogenesis (Gehlen *et al*., 2002) and afterwards (Dixit & Van Cappellen, 2002; Wu *et al*., 2017), leading to deposition of aluminosilicates in riverine systems (Presti & Michalopoulos, 2008). Diatoms might therefore be facilitating the chemical precipitation of minerals at ‘Teñidero’, as previously described for other system (Stanton *et al*., 2021), resulting in higher hydroxyaluminosilicate yields and a more intense blue coloration.

### 4.6. Conclusions

The sky-blue color of Río Celeste originates at the interface between chemistry and biology, and thus represents a textbook example of emergent behavior in environmental microbiology: by themselves, the biogeochemical processes operating in Quebrada Agria and Río Buenavista are fairly common in nature, but when put together they give rise to what could be described as a small natural wonder. Río Celeste is of relevance not only from the geobiologist’s point of view, but also for its socioeconomic impact in the Costa Rican eco-tourism industry. This unique phenomenon might however be fragile: the coloring of the river depends on just the right combination of factors, and this delicate equilibrium could very well be threatened by human activity and environmental change. Further research is therefore required in order to characterize the nature and temporal dynamics of its chemical and microbiological components, and assess its vulnerability to biotic and abiotic stressors.

## Acknowledgments

We acknowledge the support during field work from the park rangers at Tenorio Volcano National Park and the SINAC administration.

## Supplementary Figure legends

**Supplementary Table S1. Chemical composition and physicochemical properties of the three streams studied in this work.** Data taken from Castellón *et al*., 2013, DOI 10.1371/journal.pone.0075165.

**Supplementary Table S2. Sequence, abundance distribution and taxonomy of the Amplicon Sequence Variants (ASV) detected in this work.**

**Supplementary Table S3. Closest NCBI nr matches of the most abundant (more than 1000 total counts) Chloroplastic ASVs.** For each ASV, we include its best hit in the nr database to a sequence belonging to named species. In case two or more hits were tied at the best score, all of them are included.

**Supplementary Figure S1. Energy dispersive x-ray spectroscopy (EDS) of the particles shown in Figure 2.**

**Supplementary Figure S2. Fourier-transform infrared (FTIR) spectrum of the particles suspended in Río Celeste.**

**Supplementary Figure S3. Presence of diatoms around the ‘Teñidero’ confluence. a)** Relative abundance of cyanobacteria-like ASVs in the different samples. Most of the chloroplastic ASVs could be further assigned to diatoms **(Supplementary Table S3).** Sample codes are described in the legend for **Figure 3b, b)** Light microscopy image from a Río Celeste sample showing the presence of diatoms with different morphologies, **c)** Light microscopy image from a Río Celeste sample showing the presence of the dinoflagellate *Peridinium*, **d-f)** Electronic microscopy images showing details on diatom frustules.

