## Supplementary figures and images for "Out of the blue: the independent activity of sulfur-oxidizers and diatoms mediate the sudden color shift of a tropical river"

### Supplementary Figure S1.pdf

#### Río Buenavista

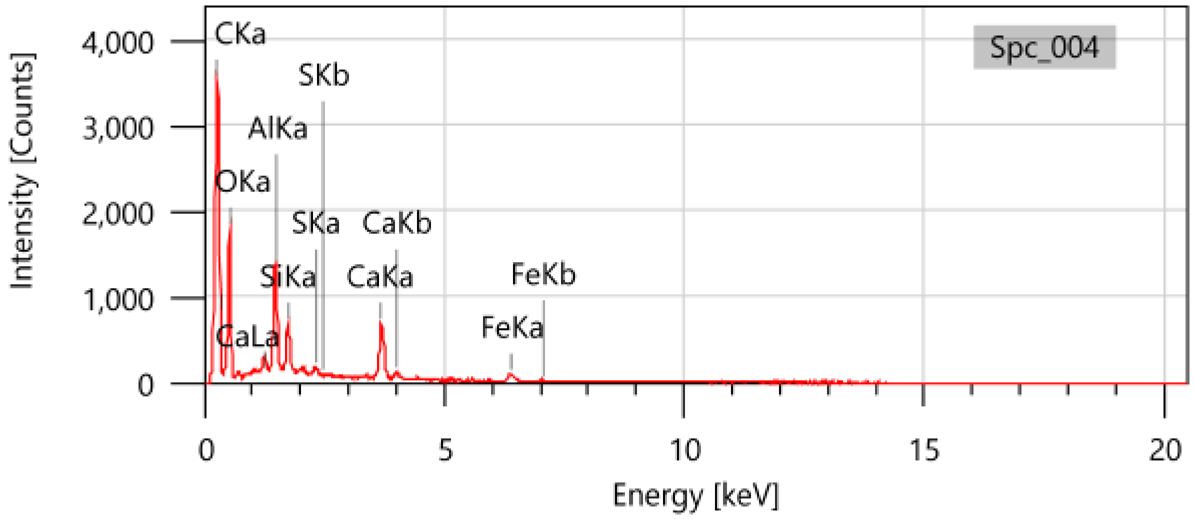


#### Río Celeste

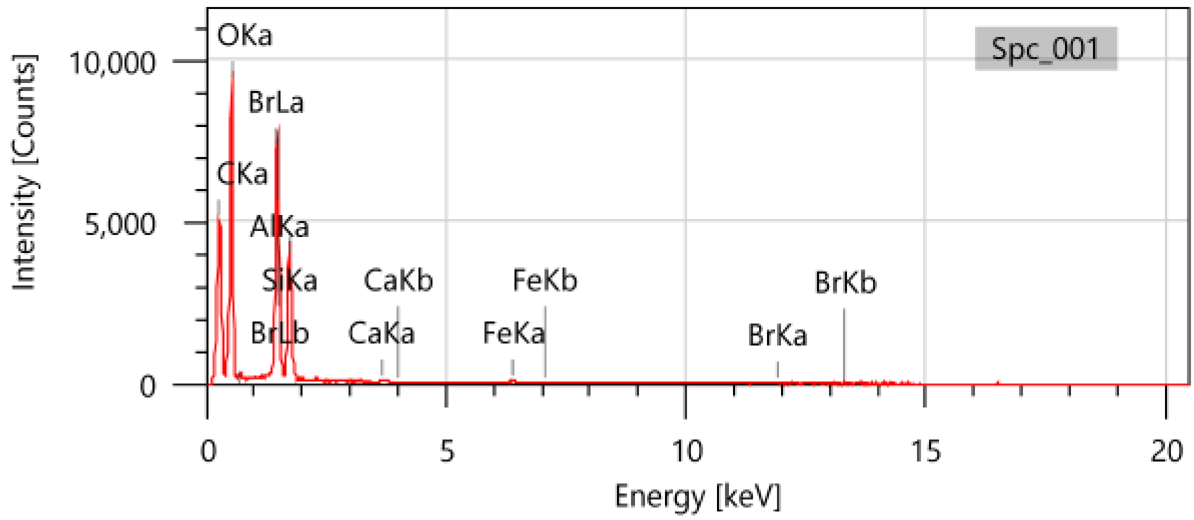


#### Quebrada Agria (neutralization experiment)

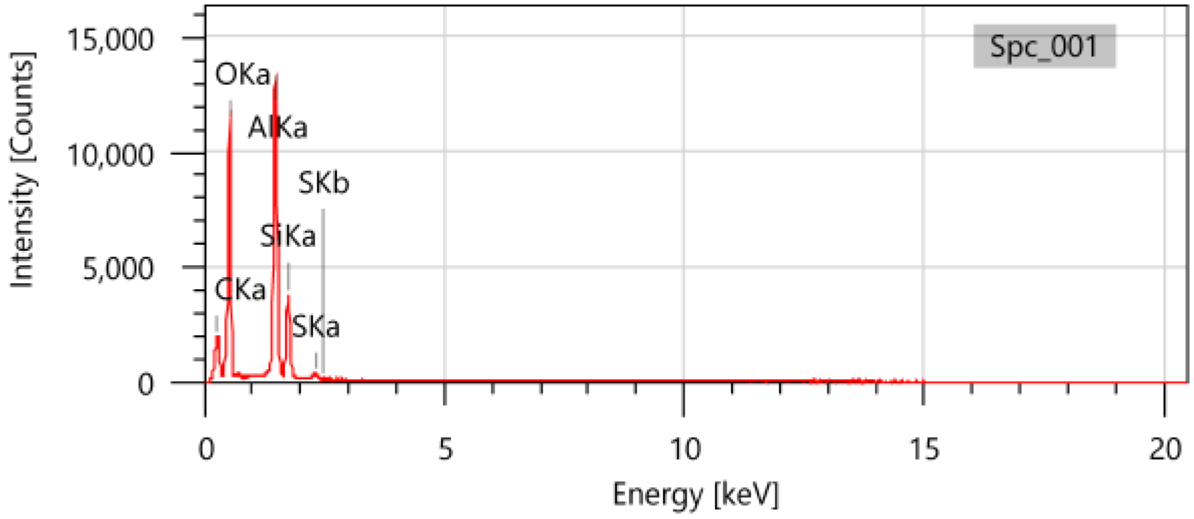

### Supplementary Figure S2.pdf

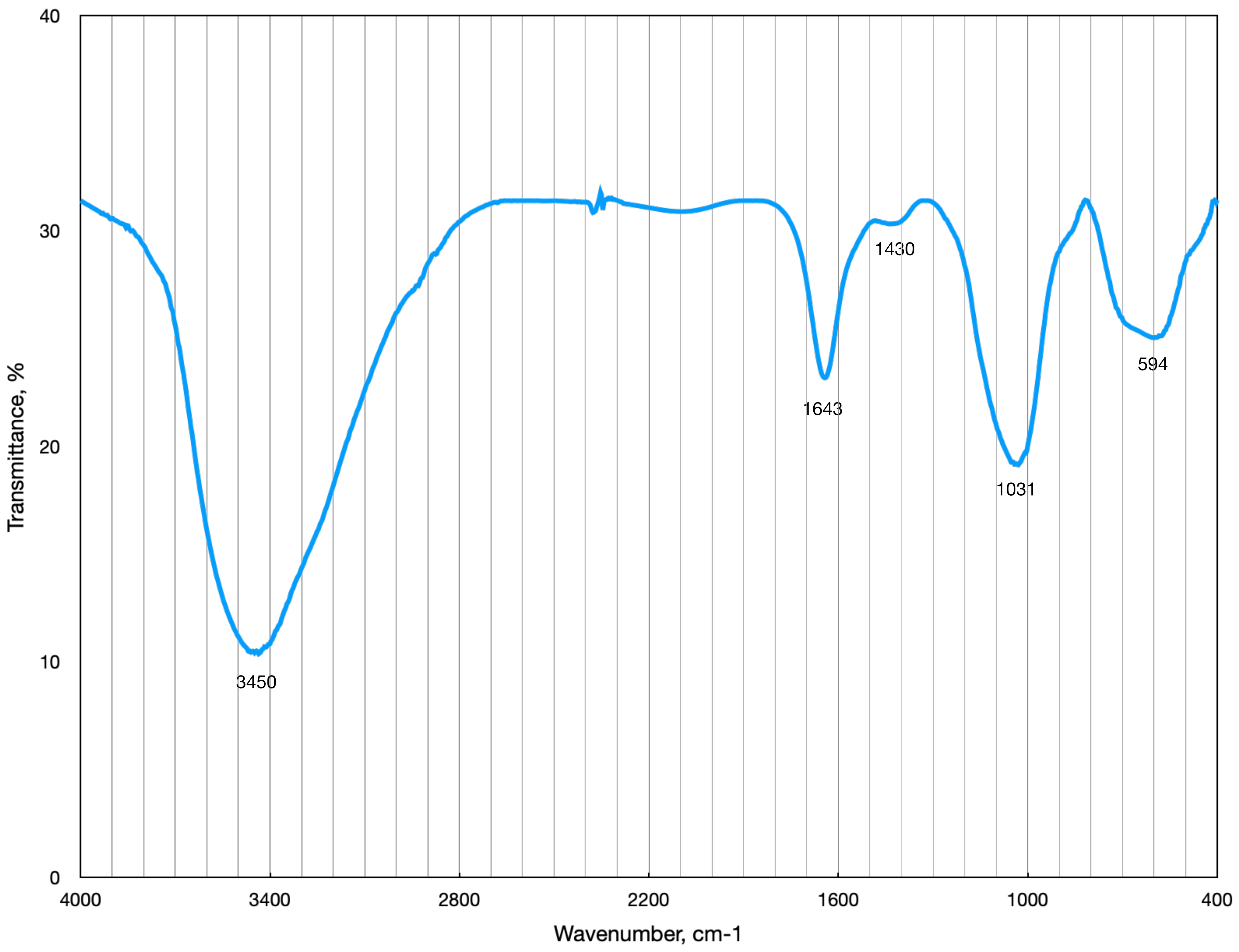

### Supplementary Figure S3.jpg

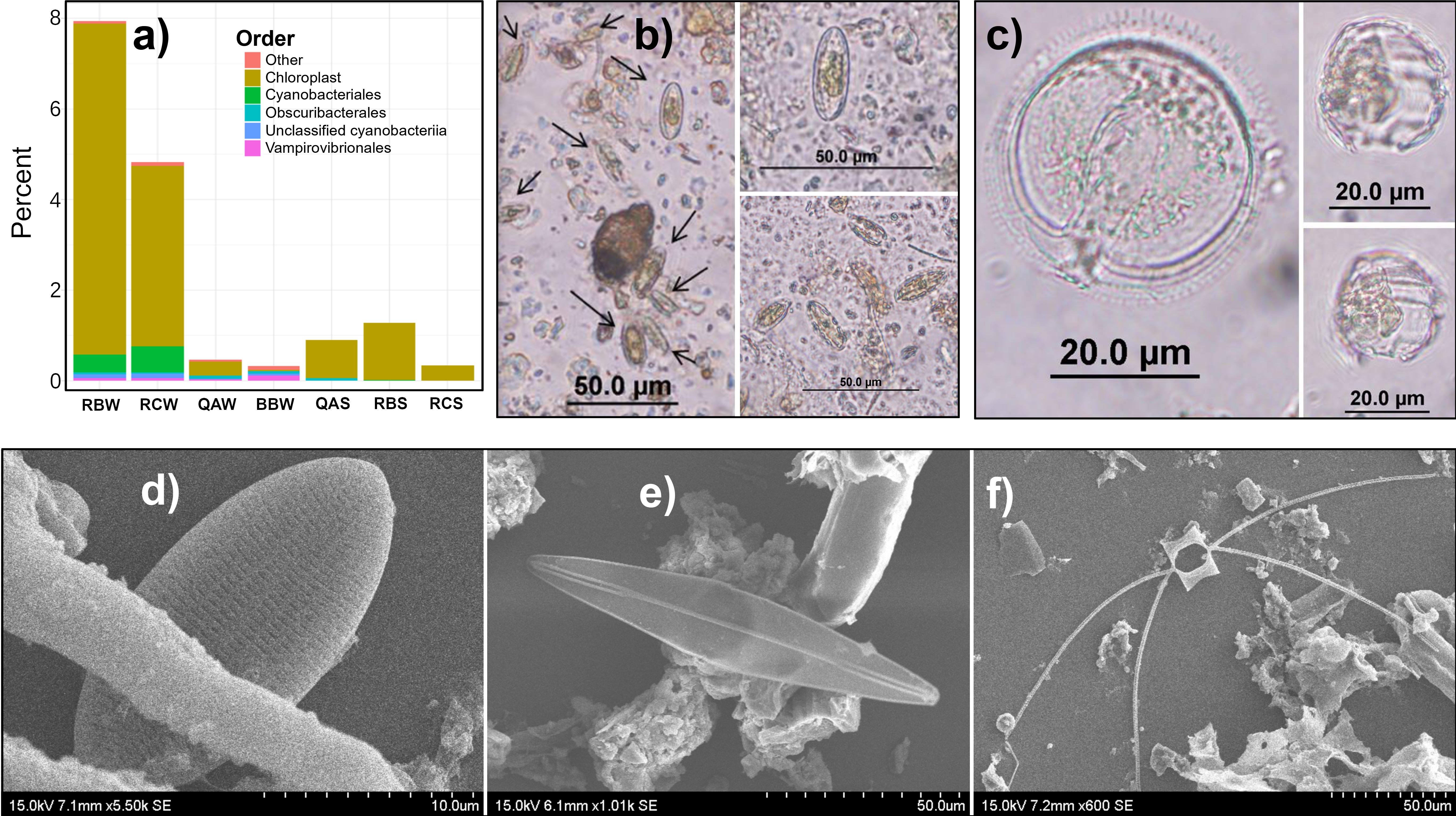
